# JUN Regulation of Injury-induced Enhancers in Schwann Cells

**DOI:** 10.1101/2022.01.31.478565

**Authors:** Raghu Ramesh, Yanti Manurung, Ki H. Ma, Todd Blakely, Seongsik Won, Eugene Wyatt, Rajeshwar Awatramani, John Svaren

## Abstract

Schwann cells play a critical role after peripheral nerve injury by clearing myelin debris, forming axon-guiding Bands of Bungner, and re-myelinating regenerating axons. Schwann cells undergo epigenomic remodeling to differentiate into a repair state that expresses unique genes, some of which are not expressed at other stages of Schwann cell development. We previously identified a set of enhancers that are activated in Schwann cells after nerve injury, and we determined if these enhancers are pre-programmed into the Schwann cell epigenome as poised enhancers prior to injury. Poised enhancers share many attributes of active enhancers, such as open chromatin, but are marked by repressive H3K27 trimethylation (H3K27me3) rather than H3K27ac. We find that most injury-induced enhancers are not marked as poised enhancers prior to injury indicating that injury-induced enhancers are not pre-programmed in the Schwann cell epigenome. Injury-induced enhancers are enriched with AP-1 binding motifs, and the c-JUN subunit of AP-1 had been shown to be critical to drive the transcriptional response of Schwann cells after injury. Using in vivo ChIP-seq analysis we find that c-JUN binds to a subset of injury-induced enhancers. To test the role of specific injury-induced enhancers, we focused on c-JUN-binding enhancers upstream of the *Sonic Hedgehog* (*Shh*) gene, which is only upregulated in repair Schwann cells compared to other stages of Schwann cell development. We confirm that c-JUN regulates these enhancers and also show that the enhancers are required for robust induction of the *Shh* gene after injury.

**Significance Statement:** The pro-regenerative actions of Schwann cells after nerve injury depends on upon profound reprogramming of the epigenome. The repair state is directed by injury-induced transcription factors, like JUN, which is uniquely required after nerve injury. In this study, we test whether the injury program is pre-programmed into the epigenome as poised enhancers and define which enhancers bind JUN. Finally, we test the roles of these enhancers by performing CRISPR-mediated deletion of JUN-bound injury enhancers in the *Sonic hedgehog* gene. While many long range enhancers drive expression of Sonic hedgehog at different developmental stages of specific tissues, these studies identify an entirely new set of enhancers that are required for Sonic hedgehog induction in Schwann cells after injury.

## Introduction

The capacity of peripheral nerve for regeneration after injury is dependent in many respects on Schwann cells (SCs). SCs undergo major injury-induced reprogramming as they are repurposed from myelin producing cells or non-myelinating cells (Remak SCs) to their nerve repair objectives (Jessen and Mirsky, 2016, 2019; Arthur-Farraj and Coleman, 2021). In the repair state, SCs clear myelin debris, secrete factors to summon macrophages, promote neuronal regeneration, create axon guidance tracks (Bungner bands), and remyelinate axons (Gomez-Sanchez et al., 2017; Jessen and Mirsky, 2019). Understanding the genesis of the repair SC is important for developing approaches to facilitate nerve regeneration, that are typically impaired in aging or in pathological conditions such as diabetic neuropathy (Painter et al., 2014; Jessen and Mirsky, 2019; Wagstaff et al., 2020; Arthur-Farraj and Coleman, 2021).

The remarkable transition of myelinating to repair SC during nerve injury is accompanied by a unique gene expression program (Nagarajan et al., 2002; Arthur-Farraj et al., 2017; Clements et al., 2017; Toma et al., 2020; Wolbert et al., 2020). SCs in injured nerve do not merely de-differentiate into a precursor cell, but rather trans-differentiate into a unique repair state which activates a gene program distinct from other stages of SC development (Jessen and Mirsky, 2016). One aspect of this unique state is defined by reliance on injury-specific transcription such as c-JUN (hereafter referred to as JUN), which is not required for Schwann cell development, but is a major early-response component for injury gene activation in SCs. A knockout of *Jun* inhibited SC repair gene induction, and caused neuronal death and reduced functional recovery after nerve injury (Arthur-Farraj et al., 2012; Fontana et al., 2012). Moreover, overexpression of JUN is sufficient to drive expression of a subset of injury genes: *Shh*, Glia Derived Neuronal Factor *(Gdnf)*, and Oligodendrocyte Transcription Factor 1 (*Olig1*) (Fazal et al., 2017). JUN is therefore a key injury-induced transcription factor, although others including STAT3 and RUNX2, may play important roles (Hung et al., 2015; Benito et al., 2017).

The novel features of the repair state extend to the use of a unique set of regulatory elements associated with the injury program. Our previous studies sought to elucidate SC reprogramming after nerve injury by assessing active enhancer regions proximal to myelin and injury genes (Hung et al., 2015). The characteristics of active enhancers include open chromatin and H3K27 acetylation (H3K27ac) (Heintzman et al., 2009; Creyghton et al., 2010; Rada-Iglesias et al., 2011; Buecker and Wysocka, 2012). We used histone mark H3K27ac to identify enhancers that become active and gain H3K27ac after injury (InjuryDB), and also active enhancers in mature SCs that lose H3K27ac after injury (ShamDB). Many ShamDB enhancers are proximal to a subset of myelin genes, which decrease in expression after injury. Conversely, many InjuryDB enhancers are proximal to injury genes which are activated after nerve injury. While a motif analysis revealed enrichment of the JUN binding motif in injury-induced enhancers (Hung et al., 2015), the scope of JUN binding in relation to injury-induced genes has not been defined.

While H3K27ac marks actively engaged enhancers, stem cell studies identified a poised enhancer state marked by open chromatin, H3K4me1, and H3K27 trimethylation (H3K27me3) instead of H3K27ac (Creyghton et al., 2010; Rada-Iglesias et al., 2011). Since H3K27me3 is a repressive mark, many of these enhancers could be activated during subsequent differentiation through active recruitment of H3K27 acetylases like CBP/p300. This raised the possibility that the injury program of Schwann cells may be pre-programmed into the epigenome through poised enhancers that could be activated after nerve injury. Our experiments test if injury gene induction is associated with poised enhancers, and we find that pioneer transcription factor activity is likely a key mechanism underpinning gene expression changes. In addition, we find that JUN binds to many injury-induced enhancers, and test the role of injury-induced enhancers in the activation of the *Sonic hedgehog* (*Shh*) gene.

## Materials and Methods

### Rat nerve injury surgery

All animal experiments were performed according to protocols approved by the University of Wisconsin, School of Veterinary Medicine (Madison, WI). Two male 4-week-old Sprague Dawley rats (Jackson Laboratory) were anesthetized under isoflurane and given an injection of 20mg/kg of ketoprofen. Under aseptic conditions, a 5 mm incision was made through the skin and muscle layers at the proximal lateral region of the femur. The sciatic nerve was exposed and cut to replicate injury. The contralateral leg received a sham operation where a 5mm incision was made just through the skin and muscle layers. The skin incision was sutured with rodent surgical staples, and the rats were caged for 8 d after surgery. The nerve tissue distal to the injury or sham site was harvested for ChIP experiments.

### Chromatin immunoprecipitation (ChIP)

Freshly dissected rat sciatic nerve was used for ChIP-seq using the MNase protocol described previously (Ma et al., 2018) with 4μg of JUN antibody (Santa Cruz SC-1694, RRID:AB_631263). Two biological replicates were performed, and samples/inputs were sequenced on a Illumina HiSeq 2500 instrument at the UW Biotechnology Center.

### Luciferase Assay

Three enhancers of mouse *Shh* as defined by the following mm10 coordinates (chr5:28557965-28558769; chr5:28567282-28568085; chr5:28630064-28630888) were amplified from mouse genomic DNA. Enhancer sites were cloned upstream of the pGL4 luciferase reporter containing the minimal E1B TATA promoter using Acc65I and BglII. RT4 Schwann cells were transfected with those enhancers and pRL-TK using TransIT-X2 (Mirus#MIR6004) and harvested for dual-luciferase assay 48hr post-transfection (n=3 per group). The luciferase assay was performed using the Dual-Luciferase Reporter assay system (Promega). A human JUN expression vector under the CMV promoter in pEZ-MO2 was purchased from GeneCopoeia (Cat. No.: EX-B0091-M02). For siRNA transfections, RT4 Schwann cells were transfected with enhancer-1 and/or 25nmol siRNAs targeting Jun (siJun-2 (IDT, DsiRNA #318163038) and siJun-3 (IDT, DsiRNA #318163041) or a negative siRNA control (IDT #51-01-14-04).

At 48 hours after transfection, total RNA was extracted and cleaned using Trizol Reagent (Invitrogen, #15596018) and RNA Clean & ConcentratorTM-5 (Zymo Research, #R1014). Jun expression was analyzed by qRT-PCR with the primers: forward GAGAGGAAGCGCATGAGGAAC, reverse CCTTTTCCGGCACTTGGAG.

### Bioinformatics

JUN ChIP-seq reads (GEO GSE190858) were mapped to the reference genome rn5 using Bowtie2 (Langmead et al., 2009; Langmead and Salzberg, 2012) to produce BAM files, and the files from the two biological replicates were combined for further analysis. BAM files were filtered for mapped reads using BamTools (Quinlan and Hall, 2010; Barnett et al., 2011) and sorted into called peaks using MACS2 (Zhang et al., 2008; Feng et al., 2012; Liu, 2014). BedTools bamCoverage generated bedgraphs of ChIP-seq samples.

Overlap analyses compared peaks from MACS2 annotated to the nearest gene, intergenic region, promoter, TSS, TES, exon, or intron via ChIPseeker (Yu et al., 2015). Pie charts showing distribution of peak sets were also generated with ChIPseeker. Heatmaps were created via EAseq (Lerdrup et al., 2016). Data processing was performed in a cloud-based manner through GalaxyBiostars (Afgan, 2018). ChIP-seq tracks were visualized using UCSC genome browser (Kent et al., 2002). Previous ChIP-seq datasets for H3K27ac (ShamDB and InjuryDB) (Hung et al., 2015), H3K27me3 and H3K4me3 (Ma et al., 2016; Ma et al., 2018) are available at GEO accession numbers: GSE63103, GSE106990 and GSE84272. RNA-seq analysis of JUN overexpression data (Fazal et al., 2017) was filtered to significant genes using the Bioconductor package DEseq (Anders and Huber, 2010). Additional expression data sets (Clements et al., 2017) were used to assess Schwann cell-specific JUN target genes. Analysis of Evolutionarily Conserved Regions in the *Shh* locus was performed using www.dcode.org (Loots and Ovcharenko, 2007).

### Generation of enhancer KO mice

The *Shh* 5’ enhancer deletion mouse line was generated by the Northwestern University Transgenic and Targeted Mutagenesis Laboratory (TTML) using CRISPR gene editing techniques. Mice were bred and housed in a specific pathogen free facility on a 12-hour light/dark cycle and fed *ad libitum* in accordance with the Northwestern University’s Institutional Animal Care and Use Committee regulations.

### gRNA identification and synthesis

The gRNA targeting the regions of interest were identified using CRISPOR online software (CRISPOR.tefor.net) (Concordet and Haeussler, 2018). These experiments utilized the AltR-Cas9 system from Integrated DNA Technologies (IDT, Coralville, Iowa), according to manufacturers’ instructions. Briefly, a sequence specific crRNA is complexed with tracrRNA (IDT, 1070532) to form an individual gRNA. Each gRNA was incubated, separately, with HiFidelity Cas9 protein (IDT, 1081064) to form individual ribonucleotide protein complexes (RNPs). Four CRISPR guide RNAs (gRNA) were designed to knockout the predicted *Shh* enhancer elements 1, 2 and 3 (Figure 4).

### Electroporation of fertilized embryos

On day 1, RNPs containing gRNA 1a (5’-ttagtccatcacctagaaag -3’) and gRNA 1b (5’-aatgcactcagataacatag-3’) were introduced into fertilized embryos as described, with minor modifications (Chen et al., 2016; Teixeira et al., 2018). Briefly, the RNP complexes were electroporated into fertilized embryos using the Electro Square Porator ECM 830 (BTX, Holliston, MA). The final concentration of reagents was 2µM of each gRNA and 4µM of HiFidelity Cas9 protein. After electroporation, the cells were cultured overnight in (blank) media at 37°C and 5% CO_2_. On day 2, RNPs containing gRNA 2a (5’-taagtgtttagcctagactc-3’) and gRNA 2b (5’-tctctgtgttggaccaccaa-3’) were electroporated into the Day1 cells using the same conditions. Electroporated cells were then transferred into pseudopregnant females as previously described.

### Genotype analysis of mice

PCR1 amplifies the WT enhancer 2 sequence and includes primers Shh_enh2_F2 5’-CTGAAAGGGCAGCAGTTACC-3’ and Shh_enh2_R2 5’-AGTAGCTGTTCACCCCACTC-3’ (expected band size 455bp). The second PCR confirms the large deletion on enhancer 1 and 2, and includes Shh_enh2_F2 5’-CTGAAAGGGCAGCAGTTACC-3’ and Shh_enh1_R1 5’-ACCATGGGACCTCAGAAGTG-3’ (expected band size 200bp). The presence of this band indicated the presence of an allele with the deletion of enhancers 1 and 2. PCR3 amplifies the WT enhancer 3 sequence and includes Shh_enh3_F15’ CGACCCTCAGCCAGTGAAG 3’and Shh_enh3_R1 5’ TCTGCCAGTTCAGTCTCTCTC 3’ (expected band size 1000bp). Finally, the deletion of enhancer 3 was confirmed with PCR4 which included primers Shh_enh3_F2 5’ TGGACAGCCCAGATAGGACT 3’and Shh_enh3_R1 5’ TCTGCCAGTTCAGTCTCTCTC 3’ (expected band size 458bp). PCR products were run on a 2% agarose gel supplemented with gel green dye (Biotium, 41005-1). A subset of F0 samples was submitted to GeneWiz for Sanger sequencing and analyzed using SnapGene software (SnapGene).

### Mouse Nerve Injury Surgery

Male and female 7-10 weeks old triple KO homozygous mice, or C57Bl6 controls, were anesthetized with isoflurane and given an injection of BupSR (sustained release) at 0.6-1.0 mg/kg SC and Carprofen (Rimadyl) at 5-10 mg/kg SC every 12 hrs for analgesia. Under aseptic conditions, a 5 mm incision was made through the skin and muscle layers at the proximal lateral region of the femur. The sciatic nerve was transected, and the contralateral leg received a sham operation where a 5mm incision was made just through the skin and muscle layers. The skin incision was sutured with rodent surgical staples, and the mice were caged for 24 hr after surgery. The nerve tissue distal to the injury or sham site was harvested for RT-qPCR analysis.

### RNA preparation, reverse transcription, and quantitative PCR

Trizol reagent was added to harvested sciatic nerve and RNA was prepared according to manufacturer’s protocol (Invitrogen cat#15596018). RNA samples were reverse transcribed on Applied Biosystems MiniAmp Thermal Cycler using QuantaBio qScript cDNA Supermix. Gene expression levels were measured by quantitative PCR on Bio-Rad CFX96 Real-Time PCR Detection System using Bio-Rad iQ SYBR Green Supermix. The following primers were used: 5’-TTAAATGCCTTGGCCATCTC-3’ and 5’-CCACGGAGTTCTCTGCTTTC-3’ for *Shh*; 5’-CGGAGTCAACGGATTTGGTCGTAT-3’ and 5’-AGCCTTCTCCATGGTGGTGAAGAC-3’ for *Gapdh*, 5’-ACTCACACACGAGAACTACCC-3’ and 5’-CCAGCTAAATCTGCTGAGGG-3’ for *Mbp*, 5’-CTGGATCGGAACCAAATGAG-3’ and 5’-GCTGAAGACCTTAGGGCAGA-3’ for *Ccl2*, 5’-GAGAGGAAGCGCATGAGGAAC-3’ and 5’-CCTTTTCCGGCACTTGGAG-3’ for cJun, and 5′-CGGGCCACTTGGAGTTAATG-3′ and 5′-TAATCTTCAGGCATATTGGAGTCACT-3′ for *Gdnf*. Levels of all transcripts were normalized to *Gapdh* levels. *Gapdh*-normalized transcript levels of injury samples were normalized again to *Gapdh*-normalized sham transcript levels using the ΔΔCT method (Livak and Schmittgen, 2001). Unpaired t tests were used for comparisons between the means of two groups (WT vs KO).

## Results

### Most injury-induced enhancers are not poised prior to injury

Previous stem cell studies defined active and poised enhancer states that share open chromatin and histone modification H3K4me1, but the key distinction is that poised enhancers are marked by H3K27me3 rather than H3K27ac (Creyghton et al., 2010). To assess whether InjuryDB enhancers are poised prior to injury, we used ChIP-seq data of H3K27me3 in rat sciatic nerve (Ma et al., 2016). Rat sciatic nerve is a good model for SC enhancer characterization since neuronal nuclei are absent and around 70-80% of peripheral nerve nuclei belong to Schwann cells. We performed an overlap analysis between the 27,277 H3K27me3 ChIP-seq peaks and previously characterized 4,311 InjuryDB enhancer peaks (Fig. 1A) (Hung et al., 2015). Surprisingly, only 45 peaks, or ∼1% of InjuryDB peaks are enriched for H3K27me3 prior to injury. Therefore, most injury-induced enhancers are not poised prior to injury.

**Figure 1.**
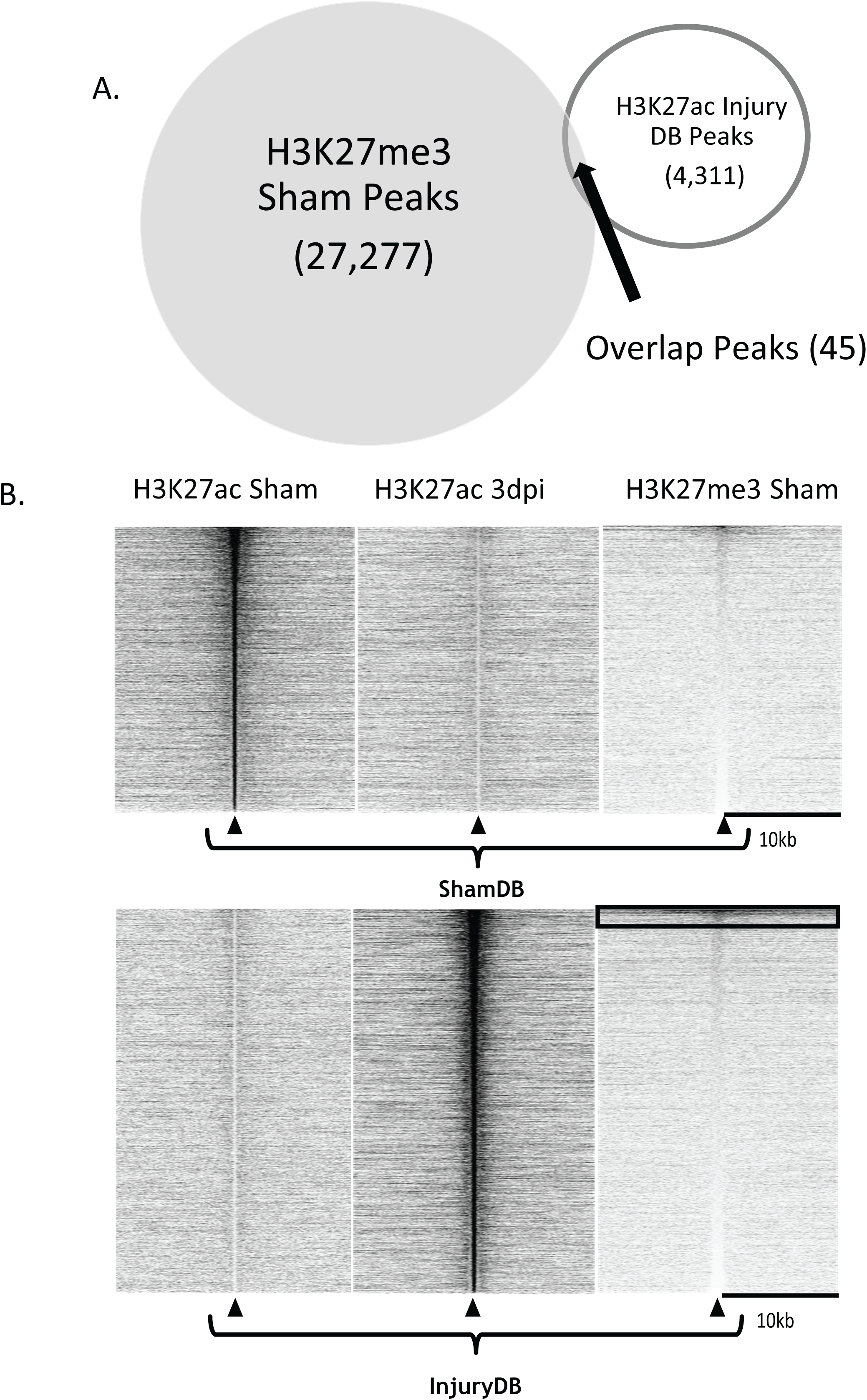
Injury-induced enhancers are not Poised. **(A)** Venn-diagram shows overlap between InjuryDB peaks and H3K27me3 sham peaks. Only 45 InjuryDB peaks have H3K27me3. **(B)** Heatmaps show distribution of H3K27ac Sham, H3K27ac Injury, and H3K27me3 Sham reads centered on ShamDB and InjuryDB peak locations, sorted in descending order by average read density. Heatmaps show enrichment of H3K27ac at ShamDB sites in the sham condition, enrichment of H3K27ac at InjuryDB sites in the injury condition, and little enrichment of H3K27me3 at InjuryDB in the sham condition. Box in H3K27me3 heatmap at InjuryDB enhancers denotes poised enhancers.

Read density plots and heatmaps were generated for H3K27ac reads in uninjured nerve (sham) and injured nerve, and compared to H3K27me3 in uninjured nerve, and the plots were centered on injury-induced enhancer peaks (Hung et al., 2015) and ShamDB peaks as a control (Fig. 1B). As expected, H3K27ac is absent on injury-induced enhancers (InjuryDB) in the sham condition, and dramatically increased after injury. Consistent with the peak overlap analysis, there is no preferential deposition of H3K27me3 at injury-induced enhancers in the sham condition, indicating that most InjuryDB enhancers are not poised.

Although poised enhancers are a small subset of InjuryDB enhancers overall, they could still be linked to important genes involved in the SC nerve injury response. We identified three poised enhancer peaks (out of 45) that are proximal to significantly induced SC injury genes (Fig. 1-1): SRY-Box Transcription Factor 2 (*Sox2*), Mesenchymal Epithelial Transition Factor (*c-Met*), and Spleen Tyrosine Kinase (*Syk*). In SCs, SOX2 is normally downregulated in myelinating cells and transgenic overexpression inhibits myelination (Le et al., 2005; Roberts et al., 2017b). SOX2 is also involved in the production of fibronectin and organizing SC migration towards the distal stump of injured nerve (Parrinello et al., 2010; Torres-Mejía et al., 2020). MET is a tyrosine kinase receptor for Hepatocyte Growth Factor (HGF) in Schwann cells and promotes peripheral nerve regeneration (Ko et al., 2018). The role of SYK is unknown in SCs, but this protein is also a tyrosine kinase that assists with diverse cellular functions such as adhesion and immune signaling (Mócsai et al., 2010).

### JUN Binds to injury-induced enhancers

In our previous study, ∼29% of injury-induced enhancers had a JUN/AP-1 binding motif (Hung et al., 2015). To assess whether InjuryDB enhancers are bound by JUN, a ChIP-seq analysis for JUN was performed at 8-days post-injury (dpi). This timepoint was chosen since some injury genes are activated over several days after injury (Ma et al., 2018), and the sustained JUN appears to be an important determinant of its activity with levels peaking at 1 week after injury (Wagstaff et al., 2020). The largest proportion of JUN ChIP-seq peaks mapped to intergenic regions (Fig. 2A). We analyzed the JUN intergenic peaks to determine whether JUN preferentially binds to the ShamDB or InjuryDB subset (Fig. 2B). As shown, JUN preferentially binds to InjuryDB enhancers with roughly 13% of those enhancers containing a JUN peak. To visualize the preferential binding of JUN on a global scale, a read density plot was generated centered on the previously defined ShamDB and InjuryDB enhancers (Fig. 2C). In line with the overlap analysis, the average read density of JUN is increased in the InjuryDB subset with minimal enrichment at ShamDB enhancers. Conversely, a similar heat map centered on the called JUN ChIP-seq peaks depicts enrichment of H3K27ac in sham and injury conditions (Fig. 2D). These plots show an increase in H3K27ac at most JUN binding sites after injury, suggesting an association between JUN binding and enhancer activation. However, there is significant binding of JUN to enhancers that were also active in the Sham condition, indicating that JUN does bind to some pre-established enhancers.

**Figure 2.**
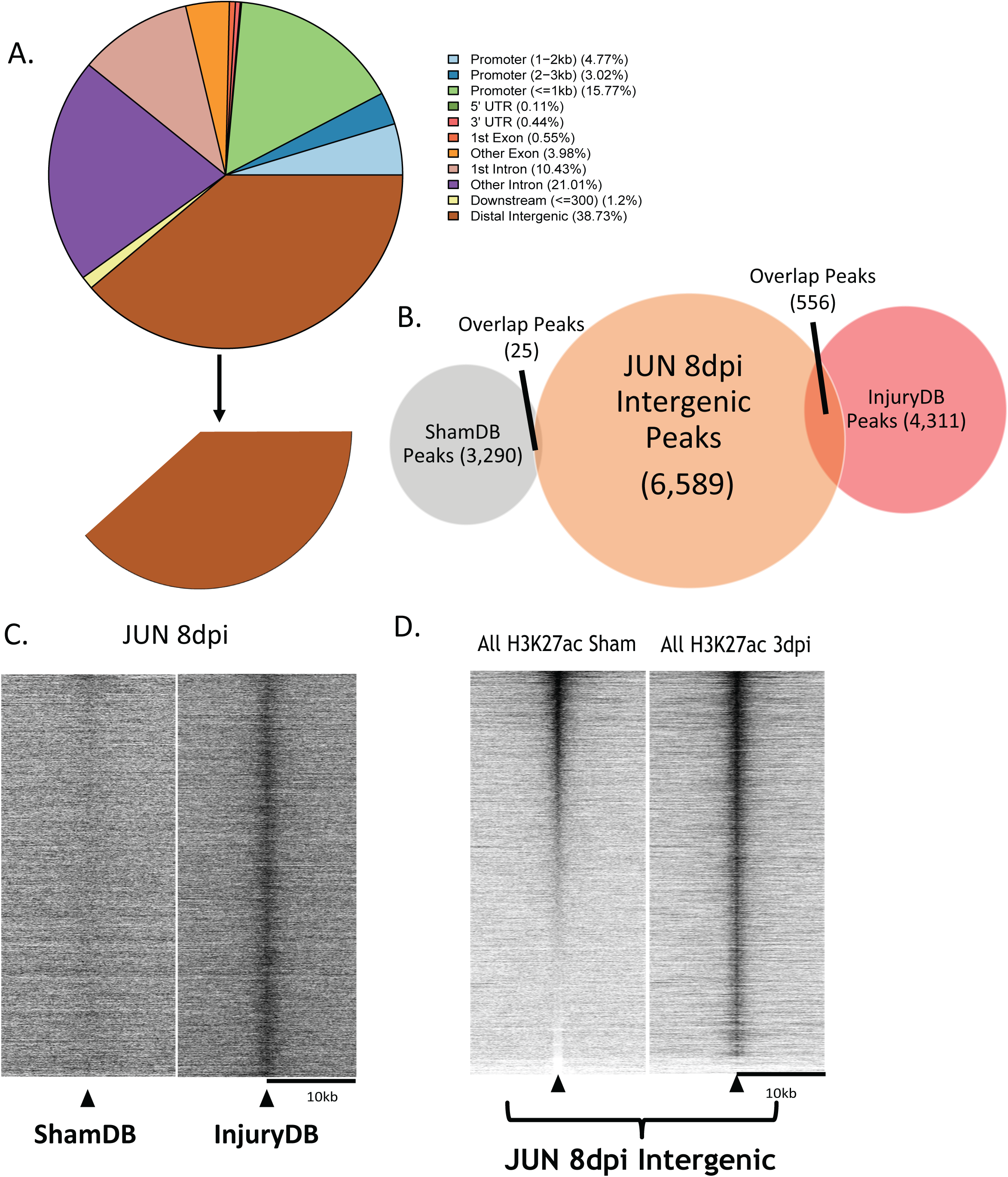
JUN Preferentially Binds at InjuryDB Enhancers After Injury. **(A)** Pie chart of annotated JUN binding peaks identified by ChIP-seq of rat peripheral nerve at 8 days post-injury. 38.52% of JUN peaks bind at distal intergenic regions. **(B)** Venn-diagram of the overlap between JUN intergenic peaks, ShamDB, and InjuryDB. Only 25 ShamDB peaks overlap with JUN intergenic peaks whereas InjuryDB peaks overlap with 556 JUN peaks. **(C)** Heatmaps show distribution of JUN ChIP-seq reads centered on the previously defined ShamDB and InjuryDB enhancer peaks. **(D)** Heatmaps show distribution of H3K27ac Sham and H3K27ac Injury ChIP-seq reads on called JUN intergenic peaks, sorted by descending average read density.

A significant proportion of JUN peaks are localized within 2 kb of transcription start sites (Fig. 2-1). It is important to note that H3K27ac is associated with actively engaged enhancers but tends to be constitutively associated with promoters (Heintzman et al., 2009), and very few promoters were identified as InjuryDB sites (Hung et al., 2015). Therefore, our H3K27ac analysis would likely have missed important JUN binding sites in promoter regions. Since not all annotated promoters are active in Schwann cells, we verified promoter localization of JUN peaks by generating a read density heat map for H3K4me3 in peripheral nerve (Ma et al., 2016), a histone mark denoting active promoters (Fig. 2-1B). A majority of JUN peak regions near transcription start sites show very strong H3K4me3 read densities centralized within a 2kb window, suggesting that most of the annotated JUN promoter peaks are indeed found at active promoter regions.

A recent publication used a transgenic overexpression approach to define genes that respond to JUN activation (Fazal et al., 2017). Using this data set, we asked how many of our 2024 promoter-localized JUN peaks overlap with genes that significantly upregulated in JUN overexpression mice (Fig. 2-1C). We compared an RNA-seq dataset of 1189 significant injury-induced genes in SCs (Clements et al., 2017) to JUN promoter peaks and found that 187 SC injury genes have JUN binding at their promoters. Out of 65 genes that are responsive to JUN and a part of the Clements et al. dataset, only 18 have active promoter JUN peaks. While JUN binding sites have been previously identified in promoter regions (Fontana et al., 2012; Norrmén et al., 2018), most of the binding sites identified in JUN-responsive genes are localized to distal enhancers in our data.

### JUN activates injury-induced *Sonic Hedgehog* enhancers

*Shh* is one of the most highly induced SC nerve injury genes and is a JUN dependent gene (Arthur-Farraj et al., 2012). *Shh* also has three distal InjuryDB enhancers that are far away from the next nearest gene, making this gene an ideal model to assess enhancer characteristics and functionality. The two proximal S*hh* enhancers are located at 65 and 76 kb upstream of the gene, and the furthest being around 180 kb upstream (Fig. 3A). None of these enhancers are poised, with the first and third enhancers also containing a JUN peak. All three enhancers have conserved AP-1 binding motifs.

**Figure 3.**
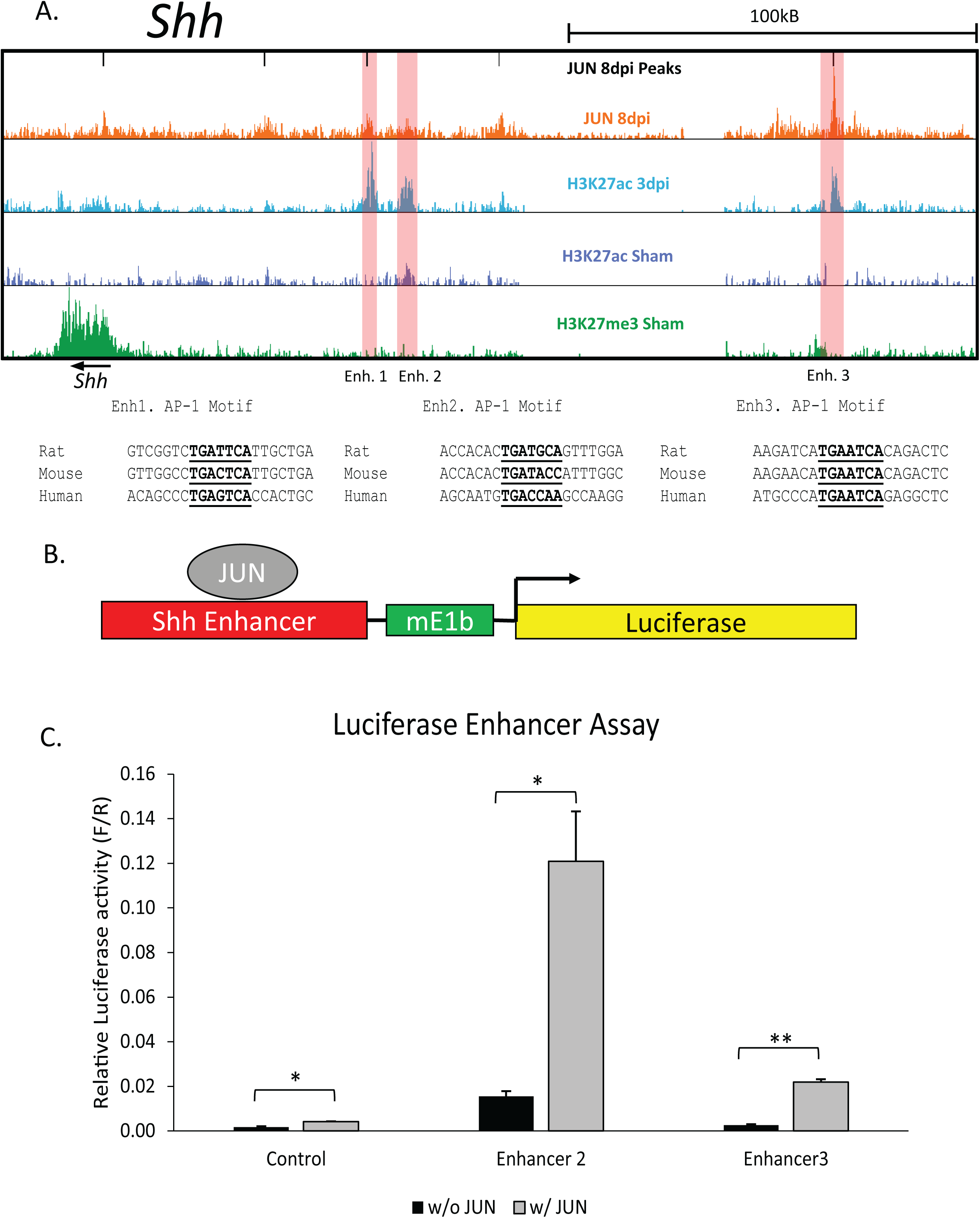
Injury-induced Enhancers upstream of *Shh* are Regulated by JUN. **(A)** ChIP-seq analysis of JUN, H3K27ac, and H3K27me3 at the *Shh* locus (n=2). The three InjuryDB enhancers proximal to *Shh* are highlighted in red. Conserved AP-1 motifs in each enhancer are shown for rat, mouse, and human. **(B)** Luciferase reporter assay for *Shh* enhancers. **(C)** Bar graph shows Shh enhancer activation after co-transfection of a JUN expression plasmid. (n=3).

**Figure 4.**
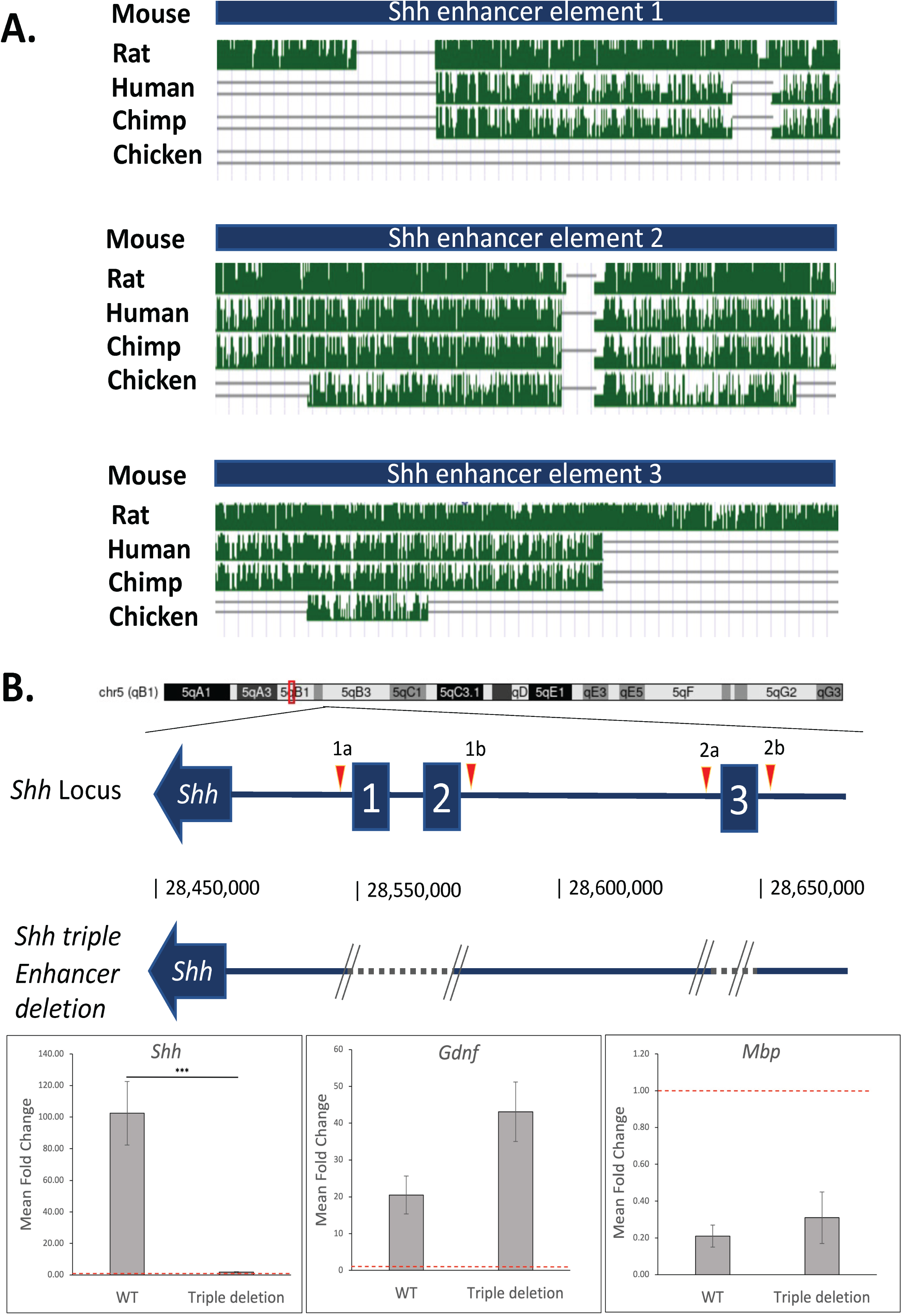
Deletion of *Shh* Enhancer Elements Abrogates *Shh* Induction after Nerve Injury. **(A)** The three injury-induced enhancers are marked among evolutionarily conserved regions upstream of the Shh gene. Red peaks are highly conserved intergenic sequences, whereas green indicates repetitive elements. Alignment of the enhancer regions using the UCSC genome browser reveled partial conservation of these regions across different mammalian species. (**B**) Top, cartoon of Shh locus showing deletion of 3 upstream *Shh* enhancer elements, that was achieved with sequential CRISPR gene editing. gRNAs targeting enhancers 1 and 2 (red triangles 1a and 1b) were electroporated into fertilized C57/Bl6 mouse embryos and cultured overnight. Embryos were electroporated with gRNA 2a and 2b (red triangles) the next day and transferred into pseudo-pregnant females. PCR analysis confirmed the deletion of the 3 enhancer elements located on Chr 5. Bottom, qRT-PCR on sciatic nerves 1-day post transection. Red dotted line indicates uninjured nerve mRNA levels. Error bars indicate s.e.m, *** indicates p<0.001, n=5 mice each.

To assess whether these enhancers can be activated by JUN *in vitro*, we created a Luciferase reporter assay in which each *Shh* enhancer was fused to a minimal E1b promoter and the luciferase gene. Co-transfection of each reporter with a JUN expression plasmid was performed in the rat RT4 Schwann cell line. *Shh* enhancer 2 was induced by JUN expression despite not having a called JUN peak *in vivo* (Fig. 3B). While Enhancer 3 was also activated by JUN, the Enhancer 1 construct had high basal activity that did not respond to JUN overexpression (not shown), so we were unable to test JUN responsiveness. However, given the high basal activity of Enhancer 1, we did transfection assays of the Enhancer 1 construct in the presence of Jun siRNA, which downregulated the activity of this enhancer (Fig. 3-1), consistent with the presence of JUN binding. In summary, JUN appears to regulate three *Shh* enhancers, consistent with the presence of conserved AP-1 motifs (Fig. 3A).

### Deletion of injury-induced enhancers reduces *Shh* expression after nerve injury

The H3K27ac-marked *Shh* enhancers are distal to the *Shh* transcription start site, leaving open the possibility that these are enhancers for neighboring genes, or that other as yet unidentified enhancers were more important for *Shh* induction. We first examined the overall conservation of these enhancers and observed that at least parts of these enhancers are conserved in mammalian species, and to a lesser extent, in birds (Fig. 4A). Next, to test the function of these enhancers in vivo, we created mice harboring a deletion of all three enhancers using CRISPR technology (Fig. 4B). To avoid a massive deletion of sequences between the first and last guide RNA (∼100kb between guide 1a, 2b), we used a sequential electroporation protocol to specifically delete the enhancer 1/2 region (∼11kb), followed by enhancer 3 region (∼1.2kb) (see Methods). After validating the deletions and breeding to homozygosity, we then tested if *Shh* was induced upon nerve transection in these mice harboring these targeted enhancer deletions. In control mice, injured nerve (1dpi) showed a ∼100-fold induction of *Shh* compared to uninjured nerve. *Gdnf* was also induced substantially in line with previous studies (Arthur-Farraj et al., 2012; Fontana et al., 2012; Arthur-Farraj et al., 2017), whereas *Mbp* was predictably reduced. In mice lacking both copies of all three enhancers, *Shh* induction was markedly reduced (Fig. 4B). *Gdnf* induction, and *Mbp* reduction was similar to controls. Our data demonstrate that these enhancers are essential for *Shh* induction.

## Discussion

During embryonic development, cell specification and differentiation are fundamental events scripted by the action of cell–specific transcription factors that determine chromatin states, and thus cell type specific gene expression. In contrast, how a fully differentiated cell reconfigures its chromatin landscape to execute a de novo, injury specific response is not well understood. SC are a fitting model to study this question, due to well defined developmental and injury states. In SC, the underlying epigenomic mechanism behind induction of injury genes is not well known. The AP-1 transcription factor JUN is required for many aspects of Schwann cell responses to injury, and both loss-of-function and gain-of-function experiments have identified a series of JUN-responsive injury genes (Arthur-Farraj et al., 2012; Fazal et al., 2017). The goal of this analysis was to elucidate the connection of JUN with injury-induced enhancers, which are marked by differential H3K27ac deposition in SCs after nerve injury (Hung et al., 2015). While recent studies have elucidated the role of several pathways regulating JUN induction (Kim et al., 2013; Kim et al., 2018; Norrmén et al., 2018), the status of injury-induced enhancers and the extent of JUN binding to such enhancers remains largely unexplored.

We first determined how many InjuryDB enhancers were poised. Poised enhancers can interact with their target promoters in a PRC2-dependent manner in differentiating ES cells (Cruz-Molina et al., 2017), but it is unclear whether SC injury induction works in this manner. We find that most injury-induced enhancers do not have H3K27me3, meaning that a majority of these enhancers are not poised. We did identify three injury genes that are proximal to poised enhancers, including *Sox2*, which is a transcription factor important for downregulation of myelin genes and SC migration during nerve injury (Le et al., 2005; Parrinello et al., 2010; Roberts et al., 2017a; Torres-Mejía et al., 2020). Although reduction of H3K27me3 through deletion of the EED subunit of PRC2 activated injury genes, most injury genes are associated with H3K27me3 in promoters/gene bodies rather than enhancers (Ma et al., 2015; Ma et al., 2016; Ma et al., 2018).

If InjuryDB enhancers are not poised or otherwise marked before nerve injury, this raises the question of how injury-induced enhancers are formed. In other systems, pioneer transcription factors have been defined as being able to bind directly to closed chromatin regions and recruit additional transcription factors to the site (Zaret and Carroll, 2011). The JUN-containing AP-1 transcription factor has many attributes of a pioneer factor. Specifically, AP-1 facilitates glucocorticoid receptor (GR) regulated transcription by binding condensed chromatin regions to allow GR binding (Biddie et al., 2011). In T-cells, AP-1 is required to open chromatin regions during T-cell activation (Yukawa et al., 2020). Interestingly, AP-1 also recruits the BAF complex, which is involved in nucleosome sliding and eviction (Vierbuchen et al., 2017). AP-1 was required to selectively open chromatin, recruit BAF, and allow for binding of cell-type specific TFs in mouse embryonic fibroblast (MEF) enhancers. If AP-1 is a pioneer factor that enables other transcription factors to bind these newly opened enhancers, this model would fit with early induction of JUN after injury (Shy et al., 1996; Parkinson et al., 2008) and the induction of other AP-1 subunits (Arthur-Farraj et al., 2017).

While there is ample evidence for the importance of JUN regulation, its binding to injury-induced enhancers had not been obtained. Here, we have obtained ChIP-seq analysis that allows us to assess how JUN interacts with injury-regulated enhancers. In our previous characterization of injury-induced enhancers, which are marked by differential H3K27ac deposition in SCs after nerve injury, we also found enrichment of the AP-1 motif in a significant proportion of these enhancers (Hung et al., 2015). Our analysis confirms JUN binding to a similar proportion of InjuryDB enhancers, although JUN also binds to pre-existing enhancers to some extent as well as several promoters of injury-induced genes, but its activity appears to be focused at enhancers.

JUN is necessary for SC repair function (Arthur-Farraj et al., 2012) because it regulates an important subset of injury genes required for a proper SC repair phenotype (Fazal et al., 2017). The importance of JUN was recently reinforced by the demonstration that the impaired regeneration caused by chronic denervation or aging could be rescued through restoring the levels of JUN (Wagstaff et al., 2020). Among the JUN target genes that have been defined, one such gene is Sonic hedgehog (*Shh*), which becomes induced within 1 day after injury, and it is sustained at higher levels for >2 weeks (Hashimoto et al., 2008; Ma et al., 2018; Wagstaff et al., 2020). *Shh* is a unique marker of the repair state in the Schwann cell lineage, as lineage tracing found no expression of *Shh* from neural crest to mature Schwann cells (Lin et al., 2015). In the nerve injury model, the role of SHH induction is not entirely clear, although it appears to have effects on axonal survival and regulation of Brain-Derived Neurotropic Factor (BDNF) (Hashimoto et al., 2008; Biddie et al., 2011). In addition, a positive feedback loop was identified where activation of *Shh* transcription by JUN leads to SHH-dependent maintenance of JUN protein levels and phosphorylation for several days after injury (Wagstaff et al., 2020). Though further analyses are required to define the interplay between these two proteins, the cooperative response of JUN and SHH hints at how JUN target genes may autoregulate JUN at a post-transcriptional level.

SHH is a potent developmental morphogen, important in many developmental contexts including limb, gut, and CNS development (Fuccillo et al., 2006; Tsukiji et al., 2014). Accordingly, there have been several efforts to identify the *Shh* enhancers that specify its induction in various tissues. In a particularly dramatic example, a highly conserved long range limb-bud specific enhancer has been identified (Lettice et al., 2003), and mutation of a single transcription factor binding site in this enhancer in snakes, is thought to account for their lack of limbs (Kvon et al., 2016). In the CNS, *Shh* expression along the rostro-caudal axis of the floor plate is driven by distinct enhancers (Anderson and Hill, 2014). Mutations in the forebrain enhancer, leads to reduced binding of SIX3 and haploinsufficiency of *SHH*, which causes holoprosencephaly (Geng et al., 2008; Jeong et al., 2008). Developmental *Shh* enhancers are exquisitely modular and spread over a megabase. These long-range enhancers are highly conserved, and direct the spatio-temporal regulation of *Shh* (Anderson and Hill, 2014; Sagai et al., 2019; Amano, 2020). However, none of these enhancers are activated in the SC injury model. Our work has characterized a novel enhancer region for *Shh*, the first such enhancers that are activated only during injury, but not in developmental states, since *Shh* is not expressed developmentally in SC (Arthur-Farraj et al., 2012; Lin et al., 2015). This enhancer does not overlap other known *Shh* enhancers, and accordingly, mice with homozygous deletions are viable and fertile with no obvious abnormalities. However, it should be noted that SLGE (*Shh* lung-gut enhancer) of *Shh* is mapped to a position between enhancers 1 and 2, and was found to drive transgenic expression to the lung/intestine (Tsukiji et al., 2014). The SLGE has also been studied as SBE6.1, which has been shown to drive CNS expression in zebrafish, mouse, and rabbit (Benabdallah et al., 2016), and its deletion in ES cells reduces expression of *Shh* in differentiated neural progenitors. While the SLGE was not marked by H3K27 acetylation in peripheral nerve, it is deleted by the Enh1/2 deletion. No developmental defects have been reported for deletion of SLGE/SBE6.1 in mouse. Interestingly, the Enh2 was designated as SBE6.2 based on sequence conservation, but it neither drove CNS expression, nor affected *Shh* levels when deleted in ES cell-derived neural progenitors (Benabdallah et al., 2016).

We show that *Shh* activation during injury is not because these enhancers are poised, but more likely due to the pioneer transcription factors, with AP-1 as prominent candidate. In the nerve injury model, SHH is tied to axonal survival and provides a neuroprotective effect by regulating Brain-Derived Neurotropic Factor (Hashimoto et al., 2008; Yamada et al., 2020). Thus, it is possible that the inability to properly activate these enhancers during aging or pathological conditions would impair nerve regeneration (Wagstaff et al., 2020). Additionally, it is conceivable that nucleotide variants in these enhancers may significantly impair nerve regeneration potential.

## Acknowledgments

The authors thank the University of Wisconsin Biotechnology Center DNA Sequencing Facility for their sequencing services and Dr. Lynn Doglio at the Northwestern Transgenic and Targeted Mutagenesis Laboratory for generation of mice. This work was supported by the National Institutes of Health: R01 NS100510 to JS, and in part by a core grant to the Waisman Center from the Eunice Kennedy Shriver National Institute of Child Health and Human Development (P50 HD105353).

## Extended Data

**Figure 1-1.**
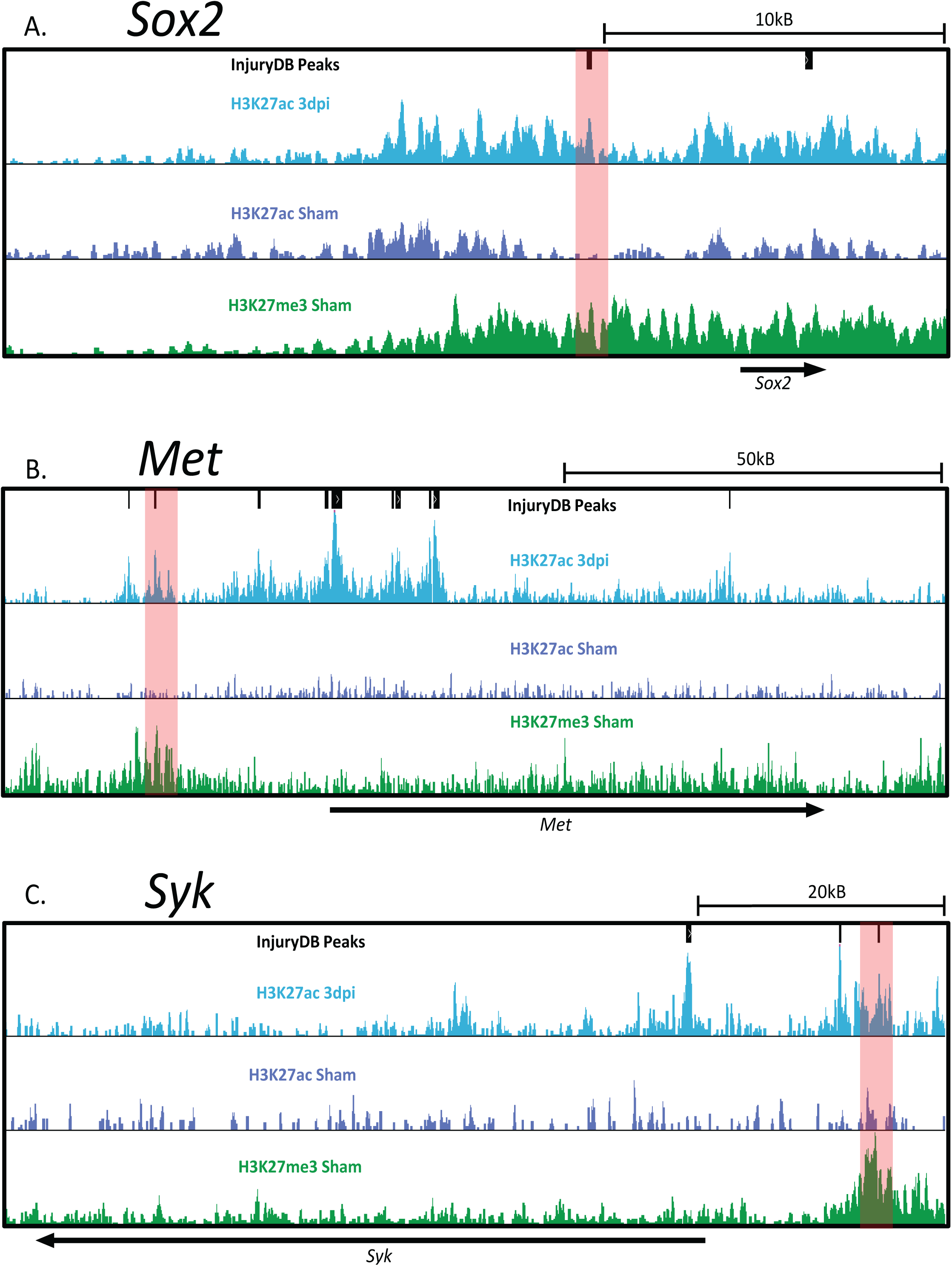
Poised Enhancers in Injury Genes. **(A-C)** ChIP-seq data on rat peripheral nerve shows the presence of H3K27me3 on injury-induced enhancers to identify 3 poised enhancers in SC injury genes (highlighted in red).: *Sox2, Met*, and *Syk*. Highlighted enhancers are conserved in mouse and human.

**Figure 2-1.**
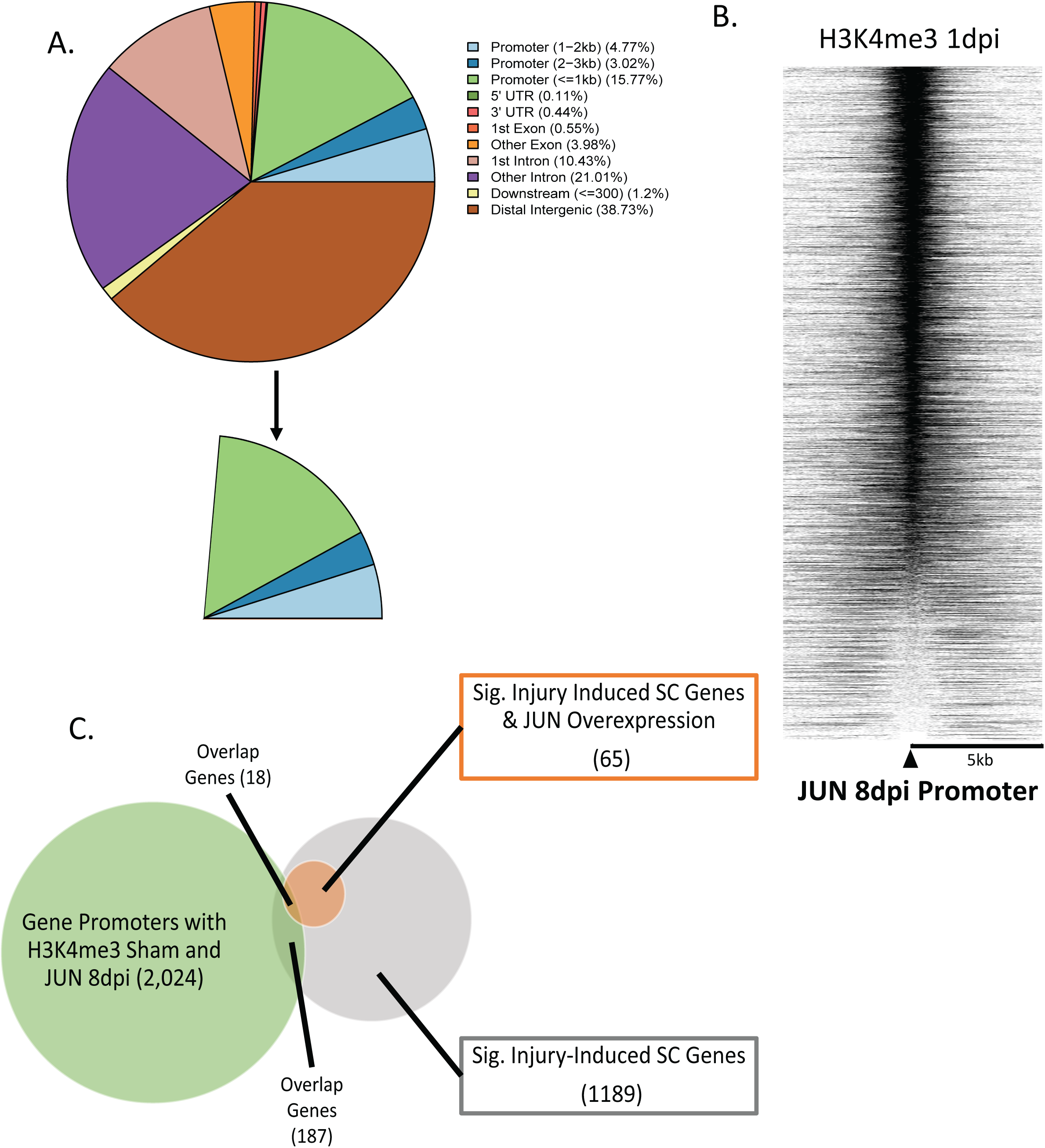
JUN Binds to Promoters of some Injury-induced genes. **(A)** Pie chart of JUN injury ChIP-seq peaks (8dpi). 22.18% of JUN peaks are found at annotated promoters. **(B)** Heatmap shows distribution of H3K4me3 reads after injury centered on JUN promoter peaks. **(C)** Venn-diagram of gene promoters that have called peaks for H3K4me3 and JUN 8dpi vs. significant JUN-dependent genes from overexpression and injury-induced SC gene datasets (Clements et al., 2017; Wagstaff et al., 2020). Only ∼8% of JUN overexpression genes overlap with gene promoters containing H3K4me3 and JUN peaks.

**Figure 3-1.**
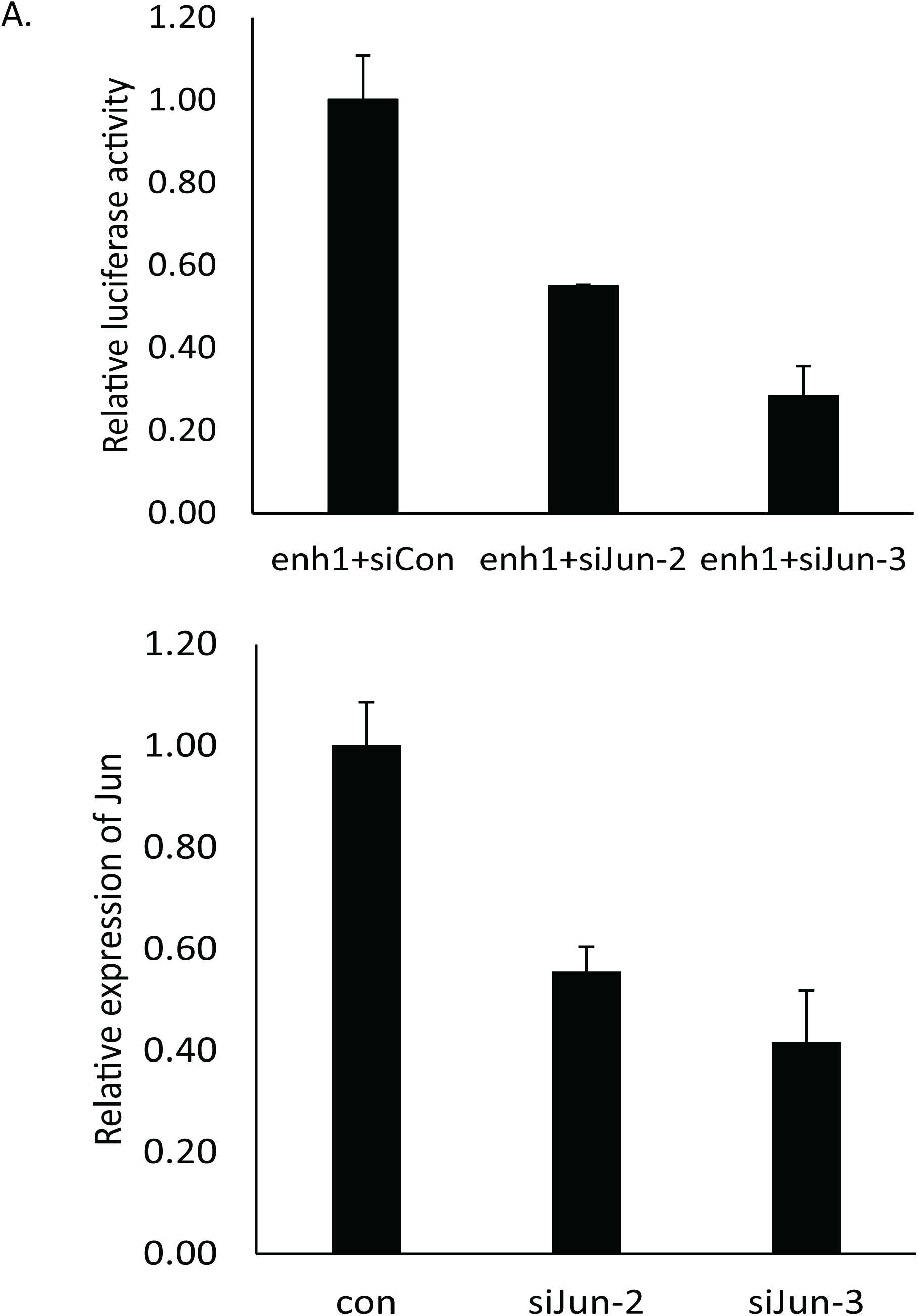
*Shh* Enhancer 1 is Regulated by JUN. **(A)** The luciferase activity of Shh Enhancer 1 was assessed after co-transfecting siRNA’s for *Jun* or a negative control siRNA into the RT4 Schwann cell line. **(B)** Independent transfections of the siRNA’s were used to evaluate their ability to downregulate endogenous Jun expression by quantitative RT-PCR.

## References

Afgan E, Andrew Taylor, JamesGoonasekera, Nuwan (2018) CloudLaunch: Discover and deploy cloud applications.

Amano T (2020) Gene regulatory landscape of the sonic hedgehog locus in embryonic development. Dev Growth Differ 62:334–342.

Anders S, Huber W (2010) Differential expression analysis for sequence count data. Genome Biol 11:R106.

Anderson E, Hill RE (2014) Long range regulation of the sonic hedgehog gene. Curr Opin Genet Dev 27:54–59.

Arthur-Farraj P, Coleman MP (2021) Lessons from Injury: How Nerve Injury Studies Reveal Basic Biological Mechanisms and Therapeutic Opportunities for Peripheral Nerve Diseases. Neurotherapeutics.

Arthur-Farraj PJ, Morgan CC, Adamowicz M, Gomez-Sanchez JA, Fazal SV, Beucher A, Razzaghi B, Mirsky R, Jessen KR, Aitman TJ (2017) Changes in the Coding and Non-coding Transcriptome and DNA Methylome that Define the Schwann Cell Repair Phenotype after Nerve Injury. Cell Rep 20:2719–2734.

Arthur-Farraj PJ, Latouche M, Wilton DK, Quintes S, Chabrol E, Banerjee A, Woodhoo A, Jenkins B, Rahman M, Turmaine M, Wicher GK, Mitter R, Greensmith L, Behrens A, Raivich G, Mirsky R, Jessen KR (2012) c-Jun reprograms Schwann cells of injured nerves to generate a repair cell essential for regeneration. Neuron 75:633–647.

Barnett DW, Garrison EK, Quinlan AR, Stromberg MP, Marth GT (2011) BamTools: A C++ API and toolkit for analyzing and managing BAM files. Bioinformatics 27:1691–1692.

Benabdallah NS, Gautier P, Hekimoglu-Balkan B, Lettice LA, Bhatia S, Bickmore WA (2016) SBE6: a novel long-range enhancer involved in driving sonic hedgehog expression in neural progenitor cells. Open Biol 6:160197.

Benito C, Davis CM, Gomez-Sanchez JA, Turmaine M, Meijer D, Poli V, Mirsky R, Jessen KR (2017) STAT3 Controls the Long-Term Survival and Phenotype of Repair Schwann Cells during Nerve Regeneration. J Neurosci 37:4255–4269.

Biddie SC, John S, Sabo PJ, Thurman RE, Johnson TA, Schiltz RL, Miranda TB, Sung MH, Trump S, Lightman SL, Vinson C, Stamatoyannopoulos JA, Hager GL (2011) Transcription factor AP1 potentiates chromatin accessibility and glucocorticoid receptor binding. Mol Cell 43:145–155.

Buecker C, Wysocka J (2012) Enhancers as information integration hubs in development: lessons from genomics. Trends Genet 28:276–284.

Chen S, Lee B, Lee AY, Modzelewski AJ, He L (2016) Highly Efficient Mouse Genome Editing by CRISPR Ribonucleoprotein Electroporation of Zygotes. J Biol Chem 291:14457–14467.

Clements MP, Byrne E, Camarillo Guerrero LF, Cattin AL, Zakka L, Ashraf A, Burden JJ, Khadayate S, Lloyd AC, Marguerat S, Parrinello S (2017) The Wound Microenvironment Reprograms Schwann Cells to Invasive Mesenchymal-like Cells to Drive Peripheral Nerve Regeneration. Neuron 96:98–114.e117.

Concordet JP, Haeussler M (2018) CRISPOR: intuitive guide selection for CRISPR/Cas9 genome editing experiments and screens. Nucleic Acids Res 46:W242–W245.

Creyghton MP, Cheng AW, Welstead GG, Kooistra T, Carey BW, Steine EJ, Hanna J, Lodato MA, Frampton GM, Sharp PA, Boyer LA, Young RA, Jaenisch R (2010) Histone H3K27ac separates active from poised enhancers and predicts developmental state. Proc Natl Acad Sci U S A 107:21931–21936.

Cruz-Molina S, Respuela P, Tebartz C, Kolovos P, Nikolic M, Fueyo R, van Ijcken WFJ, Grosveld F, Frommolt P, Bazzi H, Rada-Iglesias A (2017) PRC2 Facilitates the Regulatory Topology Required for Poised Enhancer Function during Pluripotent Stem Cell Differentiation. Cell Stem Cell 20:689–705.e689.

Fazal SV, Gomez-Sanchez JA, Wagstaff LJ, Musner N, Otto G, Janz M, Mirsky R, Jessen KR (2017) Graded elevation of c-Jun in Schwann cells in vivo: gene dosage determines effects on development, re-myelination, tumorigenesis and hypomyelination. J Neurosci 37:12297–12313.

Feng J, Liu T, Qin B, Zhang Y, Liu XS (2012) Identifying ChIP-seq enrichment using MACS. Nat Protoc 7:1728–1740.

Fontana X, Hristova M, Da Costa C, Patodia S, Thei L, Makwana M, Spencer-Dene B, Latouche M, Mirsky R, Jessen KR, Klein R, Raivich G, Behrens A (2012) c-Jun in Schwann cells promotes axonal regeneration and motoneuron survival via paracrine signaling. J Cell Biol 198:127–141.

Fuccillo M, Joyner AL, Fishell G (2006) Morphogen to mitogen: the multiple roles of hedgehog signalling in vertebrate neural development. Nat Rev Neurosci 7:772–783.

Geng X, Speirs C, Lagutin O, Inbal A, Liu W, Solnica-Krezel L, Jeong Y, Epstein DJ, Oliver G (2008) Haploinsufficiency of Six3 fails to activate Sonic hedgehog expression in the ventral forebrain and causes holoprosencephaly. Dev Cell 15:236–247.

Gomez-Sanchez JA, Pilch KS, van der Lans M, Fazal SV, Benito C, Wagstaff LJ, Mirsky R, Jessen KR (2017) After Nerve Injury, Lineage Tracing Shows That Myelin and Remak Schwann Cells Elongate Extensively and Branch to Form Repair Schwann Cells, Which Shorten Radically on Remyelination. J Neurosci 37:9086–9099.

Hashimoto M, Ishii K, Nakamura Y, Watabe K, Kohsaka S, Akazawa C (2008) Neuroprotective effect of sonic hedgehog up-regulated in Schwann cells following sciatic nerve injury. J Neurochem 107:918–927.

Heintzman ND et al. (2009) Histone modifications at human enhancers reflect global cell-type-specific gene expression. Nature 459:108–112.

Hung HA, Sun G, Keles S, Svaren J (2015) Dynamic regulation of Schwann cell enhancers after peripheral nerve injury. J Biol Chem 290:6937–6950.

Jeong Y, Leskow FC, El-Jaick K, Roessler E, Muenke M, Yocum A, Dubourg C, Li X, Geng X, Oliver G, Epstein DJ (2008) Regulation of a remote Shh forebrain enhancer by the Six3 homeoprotein. Nat Genet 40:1348–1353.

Jessen KR, Mirsky R (2016) The repair Schwann cell and its function in regenerating nerves. J Physiol 594:3521–3531.

Jessen KR, Mirsky R (2019) The Success and Failure of the Schwann Cell Response to Nerve Injury. Front Cell Neurosci 13:33.

Kent WJ, Sugnet CW, Furey TS, Roskin KM, Pringle TH, Zahler AM, Haussler D (2002) The human genome browser at UCSC. Genome Res 12:996–1006.

Kim HA, Mindos T, Parkinson DB (2013) Plastic fantastic: Schwann cells and repair of the peripheral nervous system. Stem Cells Transl Med 2:553–557.

Kim S, Maynard JC, Strickland A, Burlingame AL, Milbrandt J (2018) Schwann cell O-GlcNAcylation promotes peripheral nerve remyelination via attenuation of the AP-1 transcription factor JUN. Proc Natl Acad Sci U S A 115:8019–8024.

Ko KR, Lee J, Lee D, Nho B, Kim S (2018) Hepatocyte Growth Factor (HGF) Promotes Peripheral Nerve Regeneration by Activating Repair Schwann Cells. Sci Rep 8:8316.

Kvon EZ, Kamneva OK, Melo US, Barozzi I, Osterwalder M, Mannion BJ, Tissières V, Pickle CS, Plajzer-Frick I, Lee EA, Kato M, Garvin TH, Akiyama JA, Afzal V, Lopez-Rios J, Rubin EM, Dickel DE, Pennacchio LA, Visel A (2016) Progressive Loss of Function in a Limb Enhancer during Snake Evolution. Cell 167:633–642.e611.

Langmead B, Salzberg SL (2012) Fast gapped-read alignment with Bowtie 2. Nat Methods 9:357–359.

Langmead B, Trapnell C, Pop M, Salzberg SL (2009) Ultrafast and memory-efficient alignment of short DNA sequences to the human genome. Genome Biol 10:R25.

Le N, Nagarajan R, Wang JY, Araki T, Schmidt RE, Milbrandt J (2005) Analysis of congenital hypomyelinating Egr2Lo/Lo nerves identifies Sox2 as an inhibitor of Schwann cell differentiation and myelination. Proc Natl Acad Sci U S A 102:2596–2601.

Lerdrup M, Johansen JV, Agrawal-Singh S, Hansen K (2016) An interactive environment for agile analysis and visualization of ChIP-sequencing data. Nat Struct Mol Biol 23:349–357.

Lettice LA, Heaney SJ, Purdie LA, Li L, de Beer P, Oostra BA, Goode D, Elgar G, Hill RE, de Graaff E (2003) A long-range Shh enhancer regulates expression in the developing limb and fin and is associated with preaxial polydactyly. Hum Mol Genet 12:1725–1735.

Lin HP, Oksuz I, Hurley E, Wrabetz L, Awatramani R (2015) Microprocessor complex subunit DiGeorge syndrome critical region gene 8 (Dgcr8) is required for schwann cell myelination and myelin maintenance. J Biol Chem 290:24294–24307.

Liu T (2014) Use model-based Analysis of ChIP-Seq (MACS) to analyze short reads generated by sequencing protein-DNA interactions in embryonic stem cells. Methods Mol Biol 1150:81–95.

Livak KJ, Schmittgen TD (2001) Analysis of relative gene expression data using real-time quantitative PCR and the 2(-Delta Delta C(T)) Method. Methods 25:402–408.

Loots G, Ovcharenko I (2007) ECRbase: database of evolutionary conserved regions, promoters, and transcription factor binding sites in vertebrate genomes. Bioinformatics 23:122–124.

Ma KH, Hung HA, Svaren J (2016) Epigenomic Regulation of Schwann Cell Reprogramming in Peripheral Nerve Injury. Journal of Neuroscience 36:9135–9147.

Ma KH, Duong P, Moran JJ, Junaidi N, Svaren J (2018) Polycomb repression regulates Schwann cell proliferation and axon regeneration after nerve injury. Glia 66:2487–2502.

Ma KH, Hung HA, Srinivasan R, Xie H, Orkin SH, Svaren J (2015) Regulation of Peripheral Nerve Myelin Maintenance by Gene Repression through Polycomb Repressive Complex 2. J Neurosci 35:8640–8652.

Mócsai A, Ruland J, Tybulewicz VL (2010) The SYK tyrosine kinase: a crucial player in diverse biological functions. Nat Rev Immunol 10:387–402.

Nagarajan R, Le N, Mahoney H, Araki T, Milbrandt J (2002) Deciphering peripheral nerve myelination by using Schwann cell expression profiling. Proc Natl Acad Sci U S A 99:8998–9003.

Norrmén C, Figlia G, Pfistner P, Pereira JA, Bachofner S, Suter U (2018) mTORC1 Is Transiently Reactivated in Injured Nerves to Promote c-Jun Elevation and Schwann Cell Dedifferentiation. J Neurosci 38:4811–4828.

Painter MW, Brosius Lutz A, Cheng YC, Latremoliere A, Duong K, Miller CM, Posada S, Cobos EJ, Zhang AX, Wagers AJ, Havton LA, Barres B, Omura T, Woolf CJ (2014) Diminished Schwann cell repair responses underlie age-associated impaired axonal regeneration. Neuron 83:331–343.

Parkinson DB, Bhaskaran A, Arthur-Farraj P, Noon L, Woodhoo A, Lloyd AC, Feltri ML, Wrabetz L, Behrens A, Mirsky R, Jessen KR (2008) c-Jun is a negative regulator of myelination. Journal of Cell Biology 181:625–637.

Parrinello S, Napoli I, Ribeiro S, Wingfield Digby P, Fedorova M, Parkinson DB, Doddrell RD, Nakayama M, Adams RH, Lloyd AC (2010) EphB signaling directs peripheral nerve regeneration through Sox2-dependent Schwann cell sorting. Cell 143:145–155.

Quinlan AR, Hall IM (2010) BEDTools: a flexible suite of utilities for comparing genomic features. Bioinformatics 26:841–842.

Rada-Iglesias A, Bajpai R, Swigut T, Brugmann SA, Flynn RA, Wysocka J (2011) A unique chromatin signature uncovers early developmental enhancers in humans. Nature 470:279–283.

Roberts SL, Dun XP, Doddrell RDS, Mindos T, Drake LK, Onaitis MW, Florio F, Quattrini A, Lloyd AC, D’Antonio M, Parkinson DB (2017a) Sox2 expression in Schwann cells inhibits myelination in vivo and induces influx of macrophages to the nerve. Development 144:3114–3125.

Roberts SL, Dun XP, Doddrell RDS, Mindos T, Drake LK, Onaitis MW, Florio F, Quattrini A, Lloyd AC, D’Antonio M, Parkinson DB (2017b) Sox2 expression in Schwann cells inhibits myelination. Development 144:3114–3125.

Sagai T, Amano T, Maeno A, Ajima R, Shiroishi T (2019) SHH signaling mediated by a prechordal and brain enhancer controls forebrain organization. Proc Natl Acad Sci U S A 116:23636–23642.

Shy ME, Shi Y, Wrabetz L, Kamholz J, Scherer SS (1996) Axon-Schwann cell interactions regulate the expression of c-jun in Schwann cells. J Neurosci Res 43:511–525.

Teixeira M, Py BF, Bosc C, Laubreton D, Moutin MJ, Marvel J, Flamant F, Markossian S (2018) Electroporation of mice zygotes with dual guide RNA/Cas9 complexes for simple and efficient cloning-free genome editing. Sci Rep 8:474.

Toma JS, Karamboulas K, Carr MJ, Kolaj A, Yuzwa SA, Mahmud N, Storer MA, Kaplan DR, Miller FD (2020) Peripheral Nerve Single-Cell Analysis Identifies Mesenchymal Ligands that Promote Axonal Growth. eNeuro 7.

Torres-Mejía E, Trümbach D, Kleeberger C, Dornseifer U, Orschmann T, Bäcker T, Brenke JK, Hadian K, Wurst W, López-Schier H, Desbordes SC (2020) Sox2 controls Schwann cell self-organization through fibronectin fibrillogenesis. Sci Rep 10:1984.

Tsukiji N, Amano T, Shiroishi T (2014) A novel regulatory element for Shh expression in the lung and gut of mouse embryos. Mech Dev 131:127–136.

Vierbuchen T, Ling E, Cowley CJ, Couch CH, Wang X, Harmin DA, Roberts CWM, Greenberg ME (2017) AP-1 Transcription Factors and the BAF Complex Mediate Signal-Dependent Enhancer Selection. Mol Cell 68:1067–1082.e1012.

Wagstaff LJ, Gomez-Sanchez JA, Fazal S V., Otto GW, Kilpatrick AM, Michael K, Wong LY, Ma KH, Turmaine M, Svaren J, Gordon T, Arthur-Farraj P, Velasco-Aviles S, Cabedo H, Benito C, Mirsky R, Jessen KR (2020) Failures of nerve regeneration caused by aging or chronic denervation are rescued by restoring Schwann cell c-Jun. Elife 10:e62232.

Wolbert J, Li X, Heming M, Mausberg AK, Akkermann D, Frydrychowicz C, Fledrich R, Groeneweg L, Schulz C, Stettner M, Alonso Gonzalez N, Wiendl H, Stassart R, Meyer Zu Hörste G (2020) Redefining the heterogeneity of peripheral nerve cells in health and autoimmunity. Proc Natl Acad Sci U S A 117:9466–9476.

Yamada Y, Trakanant S, Nihara J, Kudo T, Seo K, Saeki M, Kurose M, Matsumaru D, Maeda T, Ohazama A (2020) Gli3 is a Key Factor in the Schwann Cells from Both Intact and Injured Peripheral Nerves. Neuroscience 432:229–239.

Yu G, Wang LG, He QY (2015) ChIPseeker: an R/Bioconductor package for ChIP peak annotation, comparison and visualization. Bioinformatics 31:2382–2383.

Yukawa M, Jagannathan S, Vallabh S, Kartashov AV, Chen X, Weirauch MT, Barski A (2020) AP-1 activity induced by co-stimulation is required for chromatin opening during T cell activation. J Exp Med 217:e20182009.

Zaret KS, Carroll JS (2011) Pioneer transcription factors: establishing competence for gene expression. Genes Dev 25:2227–2241.

Zhang Y, Liu T, Meyer CA, Eeckhoute J, Johnson DS, Bernstein BE, Nussbaum C, Myers RM, Brown M, Li W, Liu XS (2008) Model-based analysis of ChIP-Seq (MACS). Genome Biol 9:R137.

